# Microvilli-derived Extracellular Vesicles Govern Morphogenesis in *Drosophila* wing epithelium

**DOI:** 10.1101/2020.11.01.363697

**Authors:** Ilse Hurbain, Anne-Sophie Macé, Maryse Romao, Lucie Sengmanivong, Laurent Ruel, Renata Basto, Pascal P. Thérond, Graça Raposo, Gisela D’Angelo

## Abstract

The regulation and coordination of developmental processes involves the secretion of morphogens and membrane carriers, including extracellular vesicles, which facilitate their transport over long distance. The long-range activity of the Hedgehog morphogen is conveyed by extracellular vesicles. However, the site and the molecular basis of their biogenesis remains unknown. By combining fluorescence and electron microscopy combined with genetics and cell biology approaches, we investigated the origin and the cellular mechanisms underlying extracellular vesicle biogenesis, and their contribution to Drosophila wing disc development, exploiting Hedgehog as a long-range morphogen. We show that microvilli of Drosophila wing disc epithelium are the site of generation of small extracellular vesicles that transport Hedgehog across the tissue. This process requires the Prominin-like protein, whose activity, together with interacting cytoskeleton components and lipids, is critical for maintaining microvilli integrity and function in secretion. Our results provide the first evidence that microvilli-derived extracellular vesicles contribute to Hedgehog long-range signaling activity highlighting their physiological significance in tissue development *in vivo*.

---

Hedgehog (Hh) is a well-known long-range morphogen controlling tissue patterning and cell differentiation during embryonic development^1^. In the *Drosophila* wing imaginal disc epithelium, Hedgehog produced in posterior cells, signals across the anterior (A)/posterior (P) boundary to induce the expression of short-[i.e. *engrailed* (En), *Patched* (ptc)], and long-range [i.e. *decapentaplegic* (*dpp*)] target genes in anterior recipient cells. Different modes of carrier-mediated Hh long-range signaling have been proposed^1^, including extracellular vesicles (EVs), that arise either from fusion of multivesicular body (MVB) with the plasma membrane (exosomes), or from budding of the plasma membrane (ectosomes also termed microvesicles^2^. Evidences have been provided to suggest that both types of EVs, whose nature is tightly dependent on the site of Hh secretion (apical^3-5^ and/or basolateral secretion^6^) could mediate Hh long-range activity in *Drosophila* epithelia^4,7-9^. The apical membrane of epithelial cells is characterized by the presence of microvilli that have been shown to give rise to EVs^10,11^. However, our understanding of the mechanisms of microvilli-EV biogenesis, and their potential contribution to intercellular communication during development is presently unclear. This is mainly due to the fact that current models describing the origin of microvilli-derived EVs, focus on the relationship between the occurrence of microvilli-derived EVs and specific developmental stages^10^. These studies, which are based on ultrastructural, biochemical and pharmacological observations, reveal only part of the mechanisms of EV biogenesis and do not directly investigate the functional consequences during the developmental process *in vivo*. To understand the relationship between microvilli, EV biogenesis and their potential significance in development, we investigated the origin of EVs, the molecular mechanisms underlying their biogenesis, and their function in Hh long-range signaling using Drosophila wing imaginal disc epithelium as a paradigm.

Microvilli that are globally aligned at the apical membrane of epithelial cells, bear proteins of the Prominin family^12-14^, and have been shown, in mammalian cells, to give rise to Prominin-containing EVs^10,11^. Considering this, we examined whether the sorting of Hh to microvilli and their ability to generate EVs could be prerequisites for the EV-mediated deployment of Hh long-range signaling in the *Drosophila* wing imaginal disc (Fig. 1a). To investigate the formation of microvilli-derived EVs, we monitored the Prominin-like (PromL) protein that has been reported, upon overexpression, to distribute to apical protrusions of *Drosophila* wing imaginal discs^15^. To validate the newly generated antibodies raised against the PromL protein (methods and Supplementary Fig. 1a), we analyzed their subcellular distribution in the dorsal wing disc compartment expressing GFP-tagged PromL^15^ driven by *apterous-Gal4 driver* (*ap>PromL-GFP*) and compared to the ventral wild-type compartment within the same discs. Endogenous PromL signals massively colocalized with PromL-GFP, distributed at the uppermost apical surface above aPKC, E-cadherin (E-cad) and Dlg (apical, subapical and basolateral markers respectively), was not detected at the basolateral where PromL-GFP was previously observed^15^, and did not overlap with GFP-tagged Viking (GFP-Vkg, a collagen IV type molecule that labels the basement membrane)^16^ (Fig. 1b; Supplementary Fig 1b,c). In addition, PromL fully colocalized with Cad99c, a specific marker of microvilli of ovarian follicle and wing imaginal disc epithelial cells^17,18^ (Supplementary Fig. 1d). Overall, these results show that PromL, is associated with the apical compartment of epithelial wing disc cells, consistent with an association and distribution to microvilli.

**Figure 1.**
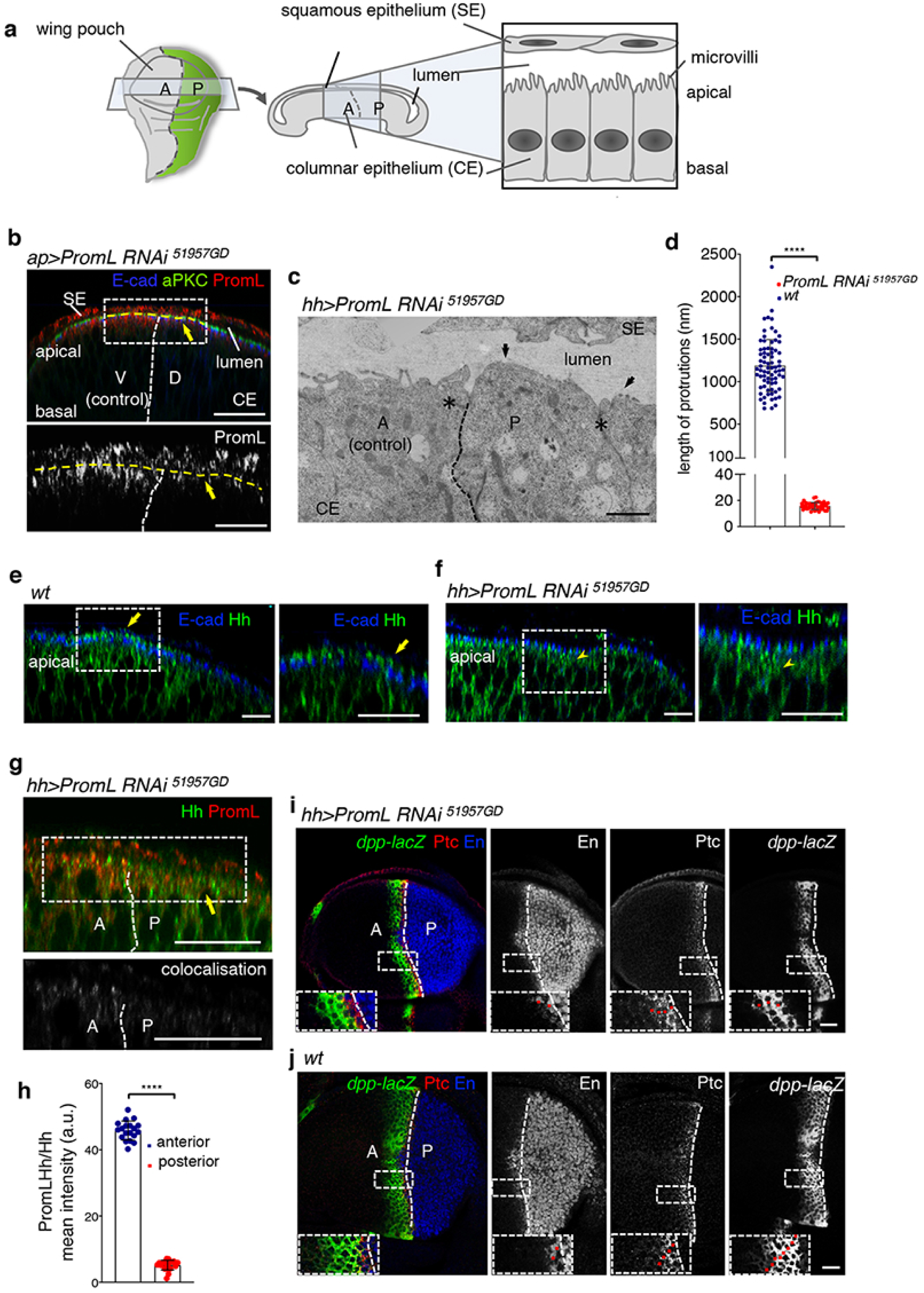
Hh long-range activity relies on microvilli integrity. **a**, Schematic of the larval wing imaginal disc epithelium and a transverse section. The disc is oriented anterior (A) to the left, posterior (P) to the right. The wing imaginal disc consists of two layers of cells with apical surfaces facing each other: the columnar epithelium (CE) of the wing disc pouch overlayered by the squamous epithelium (SE), separated by the extracellular space (lumen). Microvilli cover the apical surface of the wing imaginal disc. **b, e-g, i**,**j** Single confocal transverse and XY sections of discs of the indicated genotypes stained for PromL (red; white), aPKC (green), E-cad (blue), Hh (green), En (blue; white), Ptc (red; white) and β-gal (reflecting the expression of the reporter gene *dpp-lacZ*; green; white). **b**, Yellow arrows mark the absence of PromL in D compartment. Yellow broken line delineates the apical surface. **c**, Transmission electron micrograph of *hh>PromL RNAi* wing disc showing the absence of microvilli PromL depleted P compartment (arrows). Asterisks point to adherent junctions. **d**, Graph of microvilli length for the indicated genotypes (see Supplementary Fig. 4a). Mean ± SEM analysed by unpaired *t* test, **** *p* < 0.0001. **e, f**, Yellow arrows and arrowhead mark apical and subapical Hh distribution in *wt* and *hh>PromL RNAi* wing discs respectively. **g**, Distribution of Hh, and PromL in *wt* discs. **h**, Quantification of PromL and Hh colocalisation within the delimited region (see Supplementary information). Mean ± SEM analysed by unpaired *t* test (**** *p* < 0.0001) (n=24 uppermost apical regions, 8 discs). **i**,**j**, Confocal XY single section showing Hh target gene expression. Red dots in the insets depict the number of cells expressing En, Ptc and *dpp-*lacZ for the shown discs, but representative of all examined discs (*wt* n = 8; PromL RNAi n = 6) (see Table 1). Broken lines delimit V/D and A/P compartments. Scale bars: 1μm (**c**); 20 µm (**b, e-h**).

Next, we functionally tested the impact of PromL protein on microvilli architecture and integrity through knock down (KD) using RNA interference (RNAi) driven by *ap>* or *hedgehog-Gal4* (*hh>)* to target dorsal or posterior cells respectively. We found that in both genetic backgrounds PromL signal was specifically abrogated (Fig. 1b; Supplementary Fig. 2a). However, cell polarity (as revealed by aPKC, E-cad and Dlg staining) was not perturbed and apoptosis levels were not increased (as revealed by caspase III staining), indicating an absence of tissue stress in such condition (Fig. 1b; Supplementary Fig. 2b,c). In addition, the *wt* compartment, displayed the usual dot-like pattern of PromL reflecting a correct organization of microvilli at the apical plasma membrane (Fig. 1a,b; Supplementary Fig. 1e). We conclude that PromL specifically localised and confined to microvilli in the wing disc epithelium.

**Table 1.**
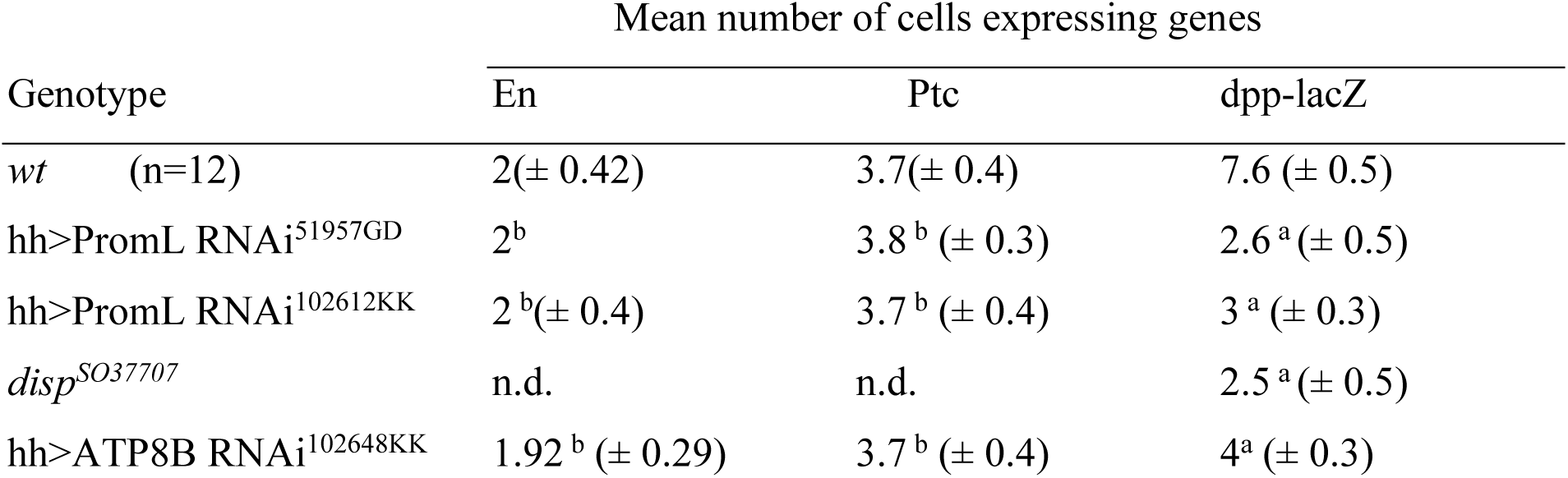
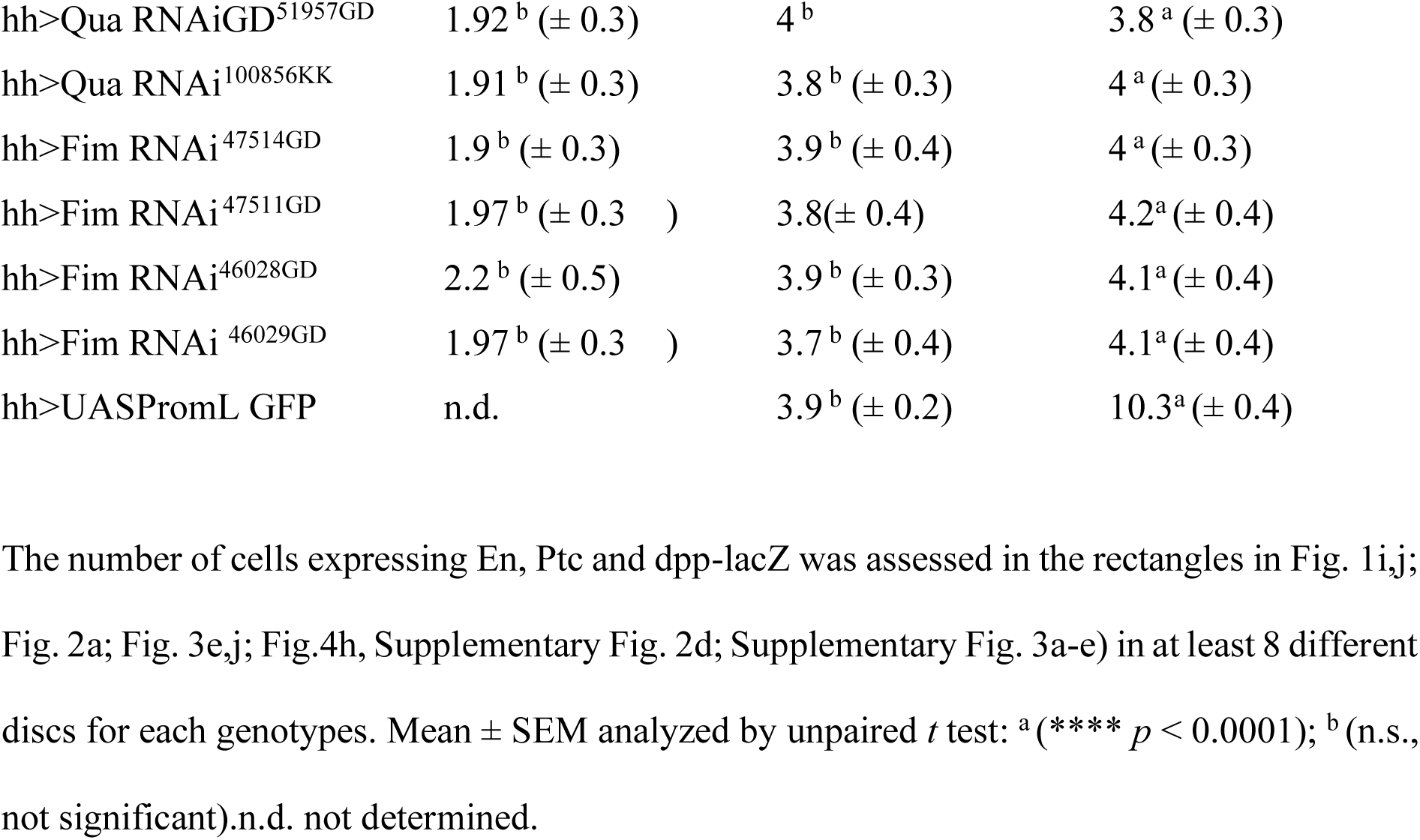
En, Ptc and ddp-lacZ expression in different genotypes.

**Figure 2.**
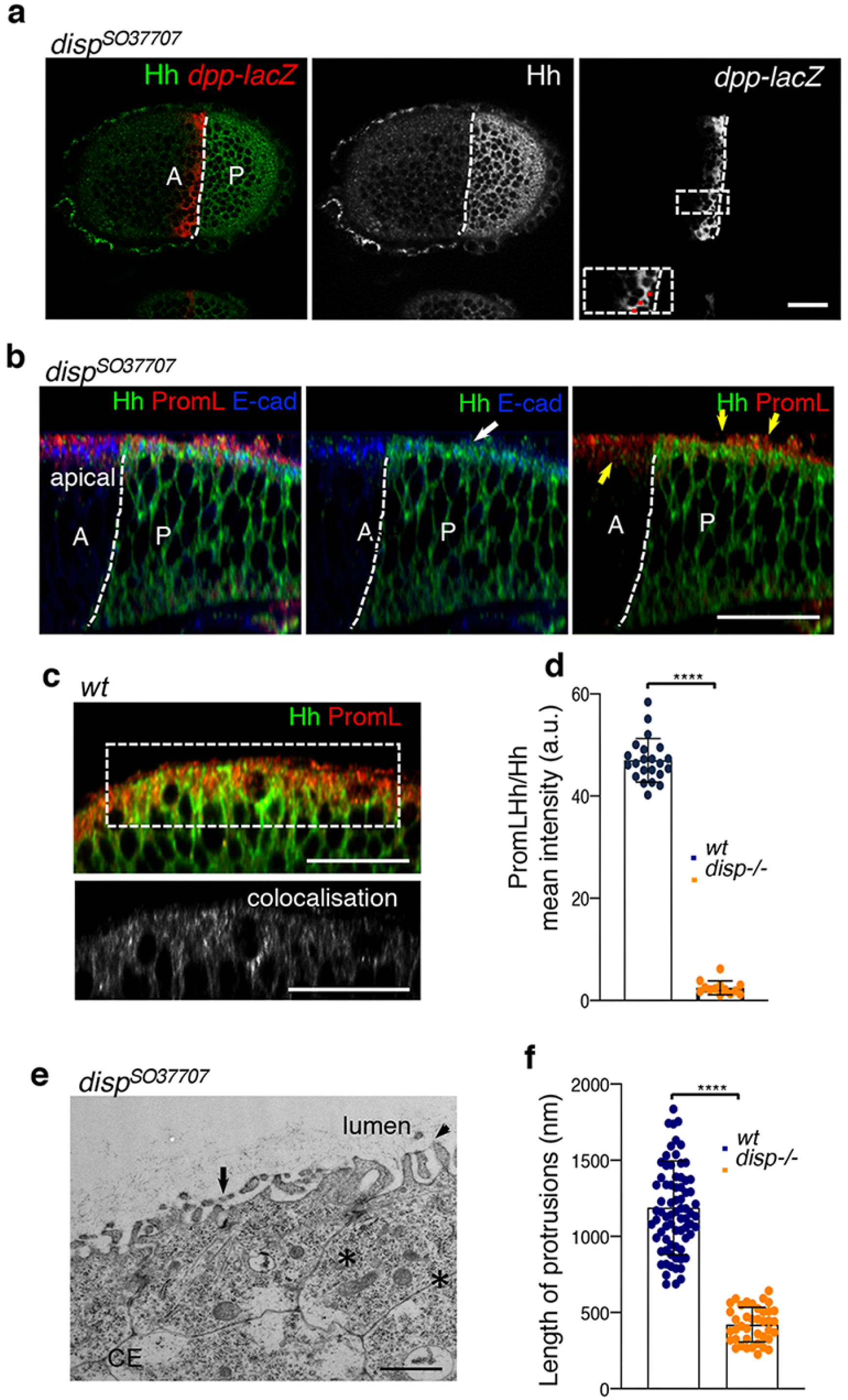
In *disp* mutant discs microvilli integrity is perturbed. **a-c**, Single confocal transverse and XY sections of discs of the indicated genotypes stained for Hh (green; white), β-gal (red; white), PromL (red; white), and E-cad (blue). **a**, Hh is retained in P compartment and *dpp-lacZ* expression is reduced to 4 cells (red dots in the inset and Table 1). **b**, Hh distributes at the apical at the level of E-cad (white arrow). Yellow arrows point to PromL staining reflecting changes in microvillus length, and compare to PromL staining in A *wt* compartment typical of the intact microvilli organization (**c**). Yellow arrows point to the similar PromL signal in A and P compartments. **c**, In *wt* discs, Hh distributed to microvilli. Bottom: shows the colocalisation of both proteins (see Supplementary information). **d**, Quantification of PromL/Hh colocalisation (see Supplementary information). Mean ± SEM analysed by unpaired *t* test (**** *p* < 0.0001) (n = 18; 6 discs). **e**, Transmission electron micrograph showing shorter microvilli (black arrows) as compare to *wt* (Fig. 4a). Asterisks depict adherent junctions. **f**, Graph reporting microvilli length. Mean ± SEM analysed by unpaired *t* test (**** *p* < .0.0001). Scale bar: 20 µm; 1μm (**e**). White dashed line delimits A/P compartments.

Upon a closer inspection of the wing imaginal disc epithelium at the ultrastructural level using transmission electron microscopy (TEM), we noticed that posterior cells of *hh>PromL RNAi* wing discs displayed abnormal microvilli or severely atrophic protrusions (with a length 98% shorter than that of microvilli from wild-type (*wt*) discs) (Fig. 1c,d; Supplementary Fig. 3a-c). In contrast, in anterior *wt* cells, microvilli length, density and morphology was similar to that of *wt* discs (Fig. 1c,d; Supplementary Fig. 4 a,b). These findings point to the requirement of PromL for the maintenance of the proper microvilli architecture and integrity in the wing disc epithelium, and are consistent with recent observation that the homologous Prominin-1 modulates microvilli architecture in mammalian cells^19^. Furthermore, our results establish that members of the Prominin protein family are key determinants of microvilli formation and integrity, putting forward the existence of a conserved function of the Prominin protein family trough evolution.

**Figure 3.**
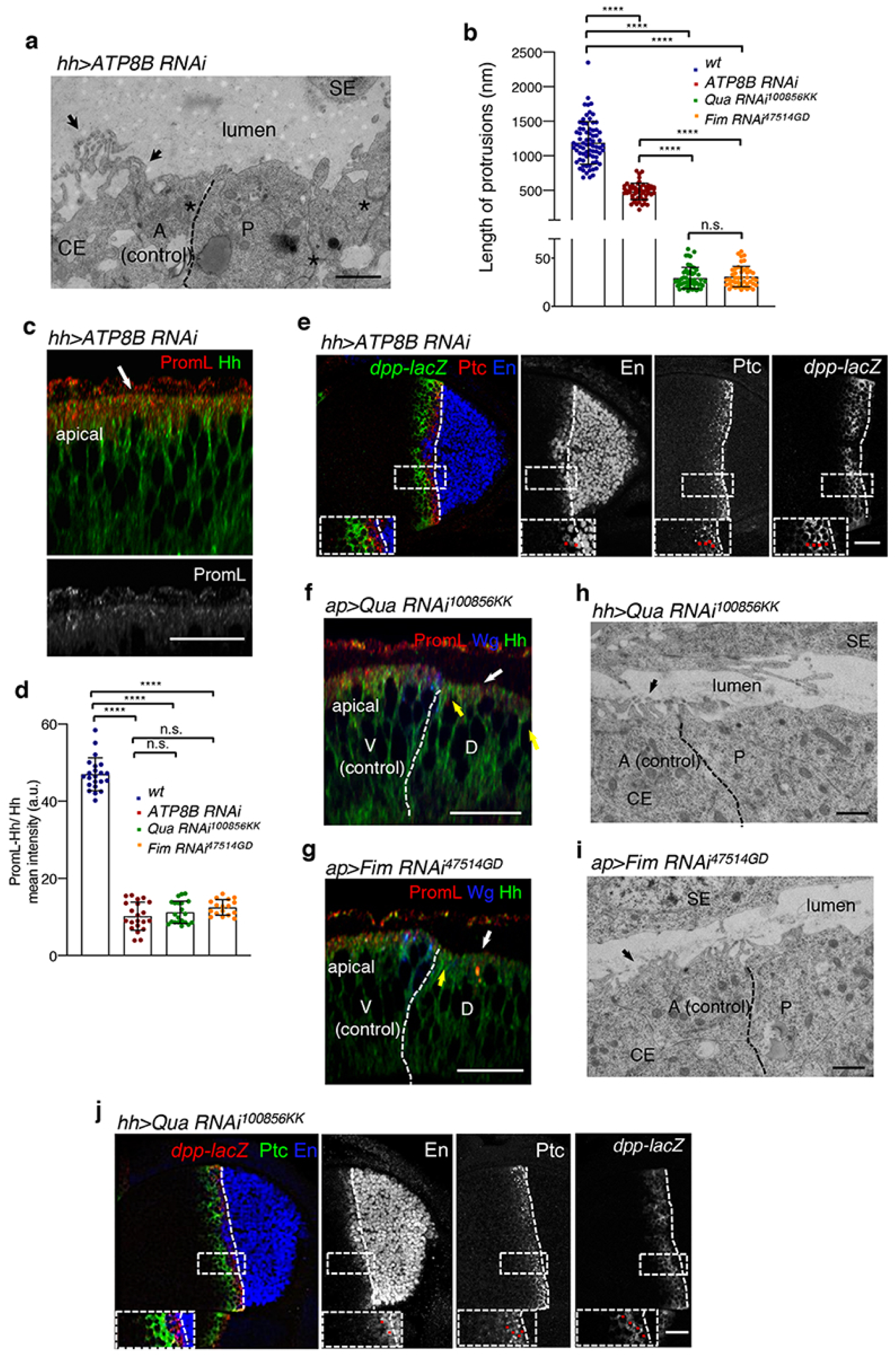
Lipids and cytoskeleton components contribute to microvilli architecture. **a**,**h**,**i**, Transmission electron micrograph of transverse sections of wing imaginal discs of the indicated genotype showing microvilli in A (black arrows), and adherent junctions (asterisk). Note that in P compartment, microvilli are reduced in size, or absent. **b**, Histogram of microvilli length. Mean ± SEM analysed by unpaired t test (**** *p* < .0.0001). n.s., not significant, *p* = 0.5230. **c**,**e**-**g**,**j**, Transverse (**c**,**f**,**g**) and XY (**e**,**j**) confocal sections of wing imaginal discs from the indicated genotypes stained for PromL (red; white), Hh (green; white), En (blue, white), Ptc (red; white), β-gal (green, white), wg (blue). **c**,**f**,**g**, Distribution of Hh and PromL for the indicated genotypes. **c**, Note absence of colocalisation of PromL and Hh in ATP8B depleted cells (white arrow). **f**,**g**, White and yellow arrows depict the absence of PromL staining and the basolateral localisation of Hh in D PromL depleted cells respectively. **d**, Quantification of PromL/Hh colocalisation in discs of the indicated genotypes. Mean ± SEM analysed by unpaired *t* test (**** *p* < 0.0001). Between 8-15 discs were analysed. **e**,**j**, Hh target gene expression in discs of the indicated genotypes. Red dots in the insets indicate the cells expressing En, Ptc and *dpp-*lacZ for the shown discs but representative of all examined discs *(ATP8B RNAi* n = 5; *Quail RNAi* n = 7; *Fim RNAi* n = 6) (see Table 1). Black (**a**,**h**,**i**) and white broken lines (**c**,**e**-**g**,**j)** delimit the A/P or V/D compartments. Scale bar: 1μm (A, B); 10 µm (C-L); 20 µm (M).

**Figure 4.**
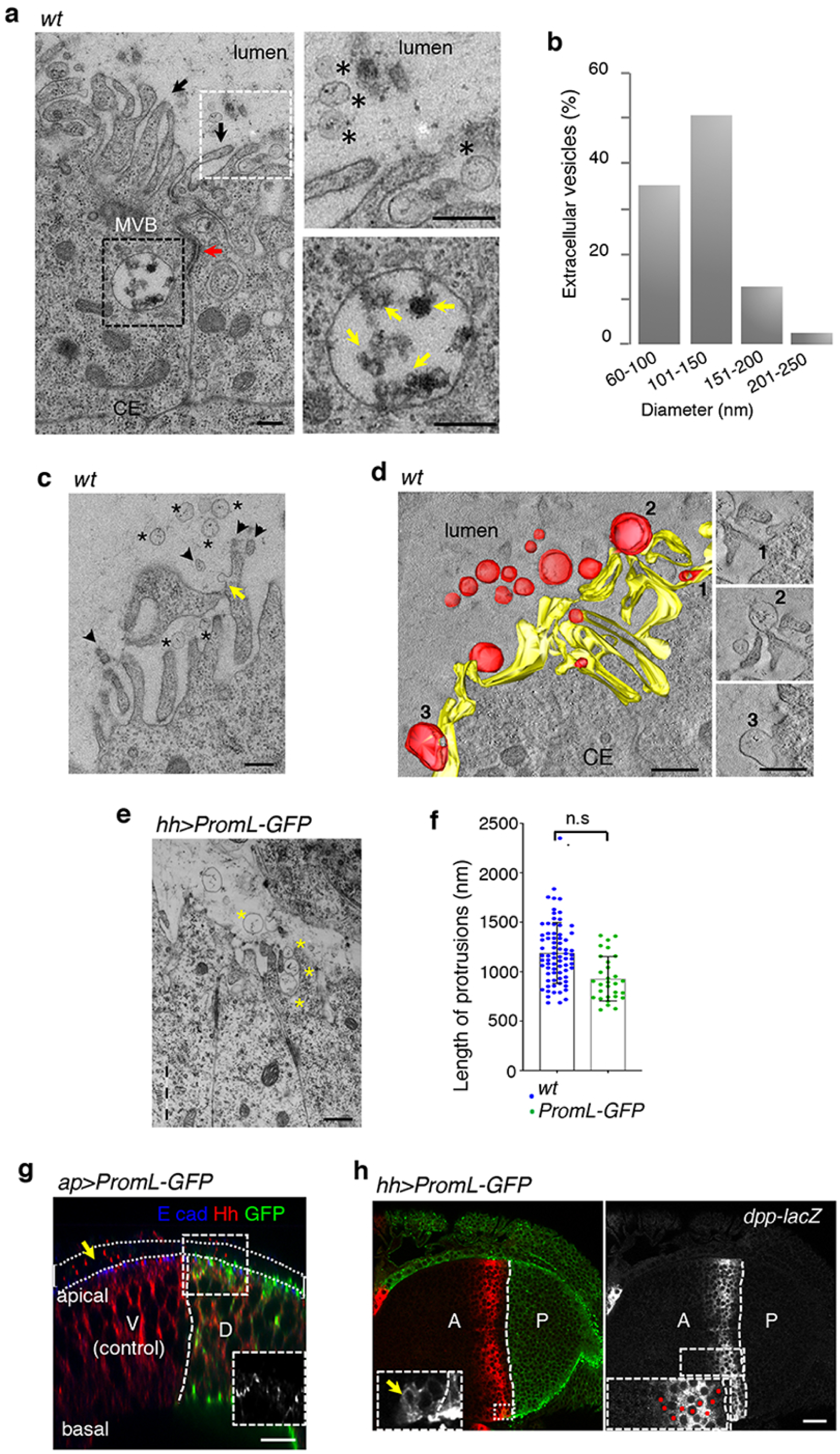
Microvilli-derived EVs control Hh long-range signaling. (**a**,**c**,**e**) Transmission electron micrograph images of a transverse section of wing imaginal discs. **a**, Microvilli (black arrows), adherent junction (red arrowhead). Top right: magnification of the white boxed region showing vesicular structures at the vicinity of the microvilli (asterisks). Bottom right shows an MVB. Yellow arrows point to ILVs. **b**, Distribution of EVs based on their diameter size (n = 32). **c**, EVs are close to microvilli (asterisks), where a bud is also visible (yellow arrow). Note that EVs are less electron dense than sections of microvilli (black arrowheads). **d**, 3D-reconstruction and modelisation of vesicles budding at microvilli. The buds (red) are still connected to the microvillar membrane (yellow) and free vesicles (red) are detected into the lumen. Right, vesicles 1-3 budding at microvilli correspond to the vesicles in (**d**). **e**, Transmission electron micrograph of *hh>PromL-GFP* disc showing EVs (yellow asterisks) into the lumen and close to microvilli (yellow asterisks). **f**, Histogram reporting the length of microvilli. Mean ± SEM analysed by unpaired *t* test (n.s., not significant), *p* = 0.4474. (**g**,**h**) *ap>PromL-GFP* (**g**) or *hh> PromL-GFP* (**h**) discs stained for E-cad (blue), Hh (red), GFP (green), and β-gal (red, white). (**g**) Hh (yellow arrow) is detected within the lumen (delimited by a white dotted line) in both D and V compartments. (**h**) XY confocal section. Left, PromL-GFP is detected in A cells (yellow arrow). Right, cells expressing *dpp-lacZ* (red dots) in the inset. Six to eight discs from each genotype were analysed, (see Table 1).Broken lines delimit the V/D (**g**) or A/P (**h**) compartments. Scale bar: 0.2 nm (**a**,**c**-**e**); 20 µm (**g**,**h**).

Having established that PromL localises to and is required for the maintenance and integrity of microvilli, we explored its relationship with Hh localization and its signaling properties. We first examined Hh distribution in *wt* and in posterior (*hh>)* or dorsal (*ap>) PromL* depleted compartments of wing imaginal discs, and found that in *wt* discs, Hh distributed at the uppermost apical surface above E-cad (Fig. 1e). No such distribution was observed in PromL mutant compartment in which microvilli were absent or misshapen (Fig. 1c), and lacking therefore a structural support for the apical distribution of Hh. Consistently, in PromL depleted cells, Hh was detected at the basolateral at the level of Dlg, and subapical to E-cad (Fig. 1f; Supplementary Fig. 2b). A subsequent quantitative analysis of the signal distribution revealed a substantial reduction of Hh staining at microvilli in posterior cells (Fig. 1g,h; methods), further supporting that the apical distribution of Hh relies on the presence of intact microvilli.

To examine the consequences of the subcellular redistribution of Hh in PromL depleted cells interferes on Hh signaling properties, we next monitored the expression of long- and short-range Hh target genes simultaneously in *hh>PromL RNAi* discs. In wing imaginal discs, *dpp-lacZ* reporter expression reflects the apical secretion and the long-range Hh signaling, whereas En and Ptc expression is attributed to Hh short-range activity mediated by its basolateral secretion^3-5^. We found that in PromL KD discs Hh long-range signaling is impaired, as evidenced by a severe decrease of *dpp-lacZ* reporter expression ranging from 60-65%. In contrast the expression of the short-range target genes, En and Ptc, remained unaltered (Fig. 1i, j; Table 1; Supplementary Fig. 2d). These results consistently establish that PromL and hence intact microvilli, are not only required for the apical distribution of Hh but also critically involved in Hh long-range signaling in the wing disc epithelium.

To further validate this hypothesis, and substantiate the requirement of microvilli integrity for Hh signaling as suggested by our observations (Fig. 1b-j), we chose to analyze the effect of Dispatched (Disp), a positive regulator of Hh secretion and trafficking^5,6,20^, on the microvillar localisation of Hh. We reasoned that in *disp* mutant discs, in which Hh is retained in producing cells, and *dpp-lacZ* expression is severely restricted (Fig. 2a; Table 1), the distribution of Hh to microvilli might also be impaired. An analysis of Hh signal showed that in *disp* mutant discs Hh consistently distributed to the apical, at the level of E-cad, but not Hh to microvilli (Fig. 2b,d), at odds with *wt* discs, in which Hh distributed to microvilli and colocalised with PromL (50% of the uppermost apical Hh signal) (Fig. 2c,d). Accordingly, long-range Hh signaling is impaired in *disp* mutants (Fig. 2a). We also noticed that PromL signal, in anterior and posterior compartment, was more punctate and irregularly spiky, pointing to a shortening of microvilli (Fig. 2b). Additional ultrastructural analysis of *disp* mutant discs indeed showed shorter and far less microvilli (66% of that of wt), with an average of 2.9 ± 1.16 microvilli per cell (Fig. 2d; Supplementary Fig. 3c). Together these experiments indicate that in addition to its role on Hh secretion and trafficking, Disp modulates microvilli architecture, reinforcing again the idea that a proper organization of apical membrane and the maintenance of microvilli architecture is critical for Hh-long-range signaling.

The maintenance of the apical plasma membrane structure relies on the asymmetrical distribution of specific phospholipids between the outer and inner leaflets of plasma membranes^21^. This is controlled by ATP8B, a putative aminophospholipid translocase, whose depletion perturbs apical membrane organization, stereocilia and microvilli integrity^22,23^. It is thus expected that abrogation of ATP8 activity can perturb microvilli organization and hinder Hh long-range signaling also in the wing imaginal disc. Consistent with this hypothesis, the qualitative and quantitative TEM analysis revealed that ATP8B depletion in posterior cells (*hh>ATPB8 RNAi)*, resulted in an alteration of microvilli number, length, and morphology, and perturbed the characteristic dot-like staining pattern of PromL indicative of the proper microvilli integrity (Fig. 3a-c; Supplementary Fig. 1e). This was accompanied by a subapical redistribution of Hh, an impediment of PromL/Hh colocalisation, and a reduction of *dpp* expression in recipient cells as a consequence of impaired long-range Hh signaling, but no effet of En and Ptc expression was observed (Fig. 3c-e; Table 1). However, ATP8B depleted cells show well developed adherent junctions, confirming their ability to polarize (Fig. 3a). Hence, loss of ATP8B activity may affect the proper distribution of PromL to microvilli, by altering the phospholipid composition of the plasma membrane, or by potentially decreasing the levels of the plasma membrane phospholipid phosphatidylinositol 4,5-bisphosphate (PIP2)^24^.

As PIP_2_, is enriched at the apical plasma membrane of epithelial cells, regulates the interaction of signaling proteins with actin-binding proteins, and promotes the bundling function of villin, it could thereby modulate the formation of actin filaments structures and cellular protrusions, including microvilli (Kumar). To test this possibility, we silenced two actin-binding proteins, Quail (Qua) (*ap>Qua RNAi*), the *Drosophila* villin-like protein^25^, or Fimbrin (Fim) *(hh>Fim RNAi)* in *Drosophila* imaginal discs. In both conditions, PromL levels were drastically reduced (Fig. 3 f,g), and microvilli, when present, were strongly defective (Fig. 3b,d,h,i; Supplementary Fig. 4c). As expected, Hh long-range signaling was impaired to similar extents in both genetic backgrounds, as confirmed by the subapical distribution of Hh and by the reduced *dpp* expression in target cells (Fig 3f,g,j; Supplementary Fig. 3; Table 1). This indicates that the disassembly of actin filaments caused profound structural alterations on the microvillar architecture and put forward the positive contribution of actin cross-linking factors in this process. Moreover, this reveals that an active interplay between PromL and cytoskeleton components contributes to the biogenesis and maintenance of microvilli that are critical for Hh-long-range signaling.

How do microvilli contribute to Hh long-range signaling? While previous work has established that microvilli give rise to Prominin-containing EVs^10,11^, their contribution to intercellular communication during development was never directly demonstrated. Considering the requirement of microvilli for Hh long-range signaling (as shown above), we reasoned that EVs could bud from microvilli and serve as a means to transport Hh to distant recipient cells. To test this assumption, we performed an EM analysis of serial sectioning of *wt* wing imaginal discs epithelial cells. We noticed the existence of small vesicular structures of 60 to 150 nm diameter, often clustered at the vicinity of microvilli, but also free within the lumen (Figure 4 a,b; Supplementary 4d; methods). We also observed that MVB with intraluminal vesicles (ILV) of smaller diameter (25-40 nm), were consistently found at close vicinity of the basolateral plasma membranes (Fig. 4a, lower right panel; Supplementary 4e), but not at the apical site, indicating that MVBs may not be prone to an apical secretion of “endosome-derived” exosomes^2^. Importantly, such vesicular structures were not detected in discs depleted for PromL, ATP8B, Qua, Fim, or *disp* mutant discs, in which microvilli are absent or atrophic, and *dpp-lacZ* expression is substantially decreased (Fig. 1c, 2e, 3a,h,i; Table 1), underscoring the correlation between the presence of microvilli, the occurrence of these secreted vesicles and Hh long-range activity. A closer inspection of the microvilli in *wt* discs, revealed the presence of buds at the microvillar membrane (Fig. 4c). Electron tomography and 3D-reconstructions of the apical plasma membrane confirmed the existence of buds still connected to the microvillar membrane, but also isolated free vesicles within the extracellular space certainly after fission (Fig. 4d, Supplementary Movies 1 and 2). Together, these results substantiate microvilli as a site of EV biogenesis and demonstrate that the active interplay between PromL, actin cross-linking proteins, and microvilli lipids is critical for the maintenance of microvilli integrity and the biogenesis of EVs. Consistent with these findings, Prominin-1 containing EVs were shown to originate from the microvilli membrane and cilia of neuroepithelial and neural progenitors cells respectively^10,26^. In agreement with our findings, actin mediate EV release from the tip of cilia^27^, and dATP8B protein is concentrated in the cilia of olfactory neuron dendrites^28^ where it could play similar roles as the one shown here in microvilli.

Given that PromL is critical for microvilli biogenesis, and that Hh distributed to microvilli where it colocalises with PromL (Fig. 1g,h; 2c,d; 3d), we reasoned that an increase of Hh long-range activity might be correlated with an increase of EV biogenesis. To test this assumption, we examined the consequences of the overexpression of PromL-GFP in Hh producing cells (*hh>PromL-GFP)*. EM analysis revealed a substantially increased of EV number (35% as compare to *wt*) in PromL-GFP discs, which resulted in a slight but not significant reduction in microvilli number and length (Fig. 4e,f; Supplementary Fig. 4b). We also found that overexpression of PromL-GFP promoted a massive Hh release into the lumen, which correlated with a significant expansion of the *dpp* expression domain (Fig. 4g,h; Table 1). These results are consistent with our hypothesis and emphasize the interdependence between PromL localisation to microvilli, EV biogenesis and Hh long-range activity.

In light of these results, we anticipate that microvilli-derived EVs, containing both Hh and PromL protein are released into the extracellular medium. To validate this hypothesis, we investigated their occurrence and their potential properties in wing discs overexpressing PromL-GFP and Hh tagged with red fluorescent protein (Hh-RFP) in Hh producing cells (*hh>Hh RFP; PromL GFP*). In a first round of experiments, we set out to investigate the distribution of Hh-GFP and Prom-GFP using immunoelectron microscopy (IEM), for a qualitative analysis. In agreement with our above observations (Fig. 1g,h; 2c,d; 3d,) both proteins localised not only to microvilli (Fig. 5a), but also on EVs (Fig. 5 b). To elucidate their dynamics and their functional roles, we investigated their properties *in vivo*. Live imaging of *hh>Hh RFP; PromL GFP* discs revealed that the majority (80%) of EVs released by the posterior compartment was positive for Hh-RFP and Prom-GFP, whereas the others only displayed Hh-RFP (Fig. 5c-e). These EVs were not static but engaged in a dynamic movement and trafficked associated to one another within the extracellular space (Fig. 5f,g; Supplementary Fig. 5a; Supplementary Movies 3). These EVs are very likely the extracellular carriers of Hh, facilitating thereby its long-range activity. These results are consistent with our hypothesis that microvilli of the wing disc epithelium are the site of generation of small EVs, that are critically involved in the transport of Hedgehog across to facilitate its long-range activity.

**Figure 5.**
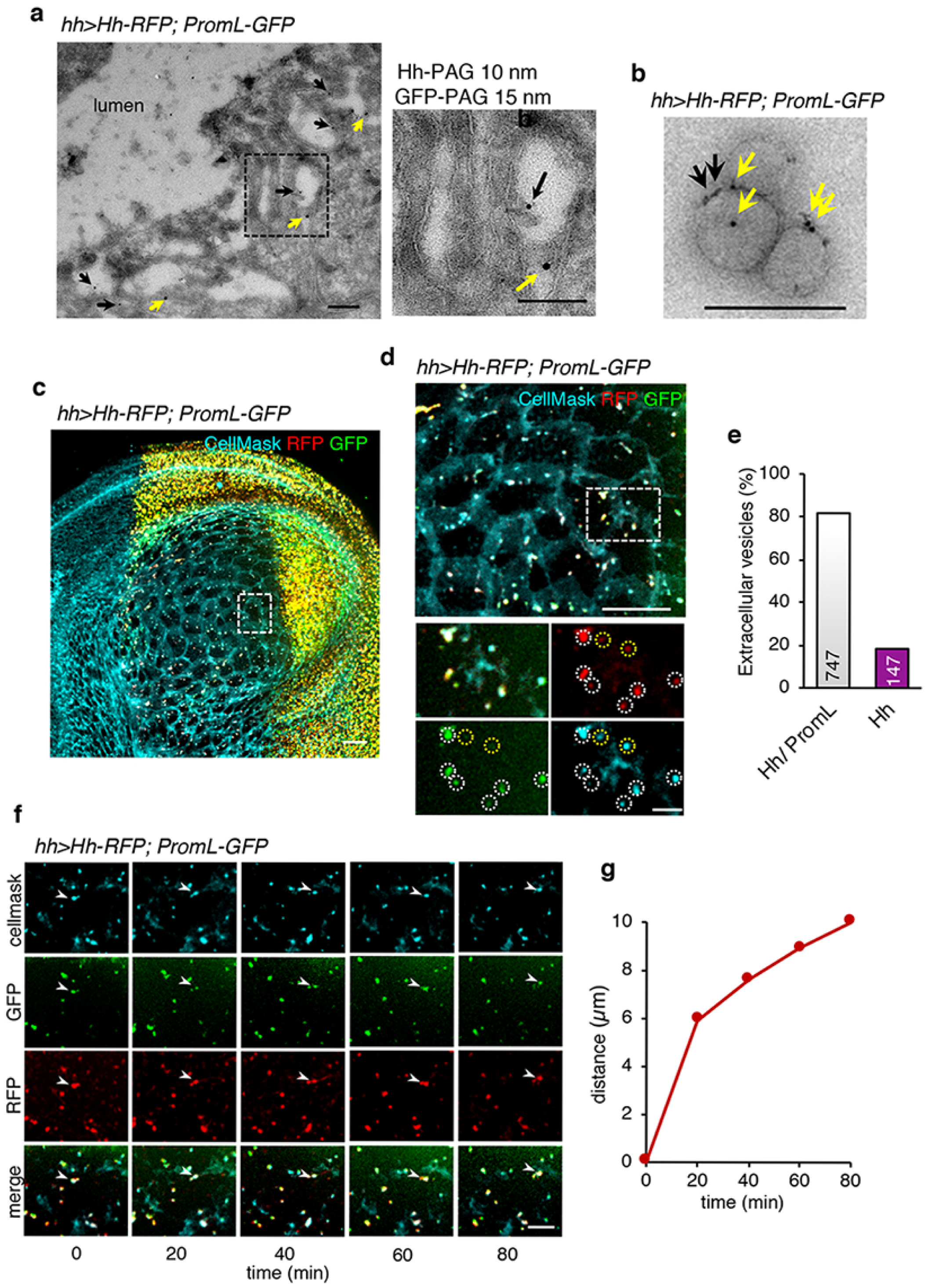
Hh-PromL-EVs move into the extracellular field. **a**, Ultrathin cryosection of *hh>Hh RFP; PromL-GFP* disc immunolabeled for Hh (PAG 10; black arrows) and GFP (PAG 15, yellow arrows) showing both proteins at microvilli. **b**, EVs detected into the lumen are positive for Hh (black arrow), and PromL (yellow arrow). **c**, Maximum intensity *z*-projection of 167 slices spaced 1 µm apart of *hh> Hh-RFP; PromL-GFP* disc recorded every 3 min. Plasma membrane was stained with Cellmask (Cyan). **d**, Magnified area of *z*-projections of 167 plans, spaced 1 µm apart, displaying a XY view of the anterior compartment of *hh> Hh-RFP; PromL-GFP* disc recorded every 3 min for 90 min. Bottom, magnification of the inset showing examples of Hh/PromL-containing EVs (white circles) or Hh-containing EVs (yellow circles). **e**, Histogram of Hh/PromL-EVs or Hh-EVs counted in the anterior compartment at different times and in different regions, (n = 894). **f**, Frames from Movie 2 of *hh>Hh-RFP; PromL-GFP* disc at the indicated time-points. **g**, The travel distance of Hh/PromL-EVs (white arrowheads) gradually increased with time and is of 10µm in 80 min for the shown vesicle. The manual tracking of EVs was performed in the boxed region in (**c**). Scale bar: 0.2 µm (**a**,**b**); 20 µm (**c**,**f**); 5 µm (**d**,**f**).

Understanding the origin and function of EVs *in vivo* during development has so far been limited especially because of a shortage of methods suitable to demonstrate causality *in vivo*^10,11^. Taking advantage of complementary electron and fluorescence microscopy, live imaging, cell biology and genetic approaches, our experiments unveil that EVs provide a means for exchanging signaling cues between cells at distance. Here, we find that PromL, by interaction with lipids and cytoskeleton components, is a key determinant of microvilli formation *in vivo*, whose integrity is critical for the biogenesis of EVs and their signaling role (Fig. 1; Fig. 3). Finally, we uncover that microvilli are the preferential site for the generation of small EVs conveying Hh, - different form exosomes-, revealing the existence of a new potential mechanism mediating Hh long-range signaling in the *Drosophila* wing imaginal disc epithelium, and unveil their physiological significance in tissue development *in vivo*.

Considering that the morphogen Hh controls a variety of conserved functions in invertebrates and vertebrates, and that the dysregulation of the Hh pathway promotes developmental defects and contributes to several cancer types, it will be of high interest to investigate the contribution of the EV-mediated Hh signaling described in this study to such physiological and pathological processes.

## Acknowledgments

We thank A. Plessis and Ph. Vernier for insightful discussions. C. Dahmann, X., D. Godt, the Developmental Studies Hybridoma Bank, the Vienna *Drosophila* RNAi Center, *Drosophila* Genetics Resource Center, and Bloomington stock centers for reagents. Vincent Fraisier from the Imaging platform UMR144; The Cell and Tissue Imaging (PICT-IBISA), Institut Curie, member of the French National Research Infrastructure France-BioImaging (ANR10-INBS-04). This work was supported by the French Government (National Research Agency, ANR) through the Investments for the Future LABEX SIGNALIFE (ANR-11-LABX-0028-01), LabEx CelTisPhyBio (ANR-11-LABX-0038), ANR (ANR-15-CE13-0002-01), Ligue Contre le Cancer ‘‘Equipe labellisée 2016’ to P.P.T, and by the Fondation pour la Recherche Médicale: DEQ201110421324.

## Authors contributions

Conceptualization: P.P.T, G.R, and G.D.; Methodology: I.H, M.R, L.S., L.R., and G.D.; Software: A.S.M.; Formal analysis: I.H, A.S.M, and G.D.; Investigation: I.H., M.R., L.R., G.D.; Resources: R.B., P.P.T., G.R.; Writing - Original Draft: G.D.; Writing - Review & Editing: I.H., R.B., P.P.T., G.R. and G.D.; Supervision: G.D.; Funding acquisition G.R. and P.P.T.

## Data and materials availability

all data is available in the manuscript or the supplementary information.

---

**Supplementary Figure S1.**
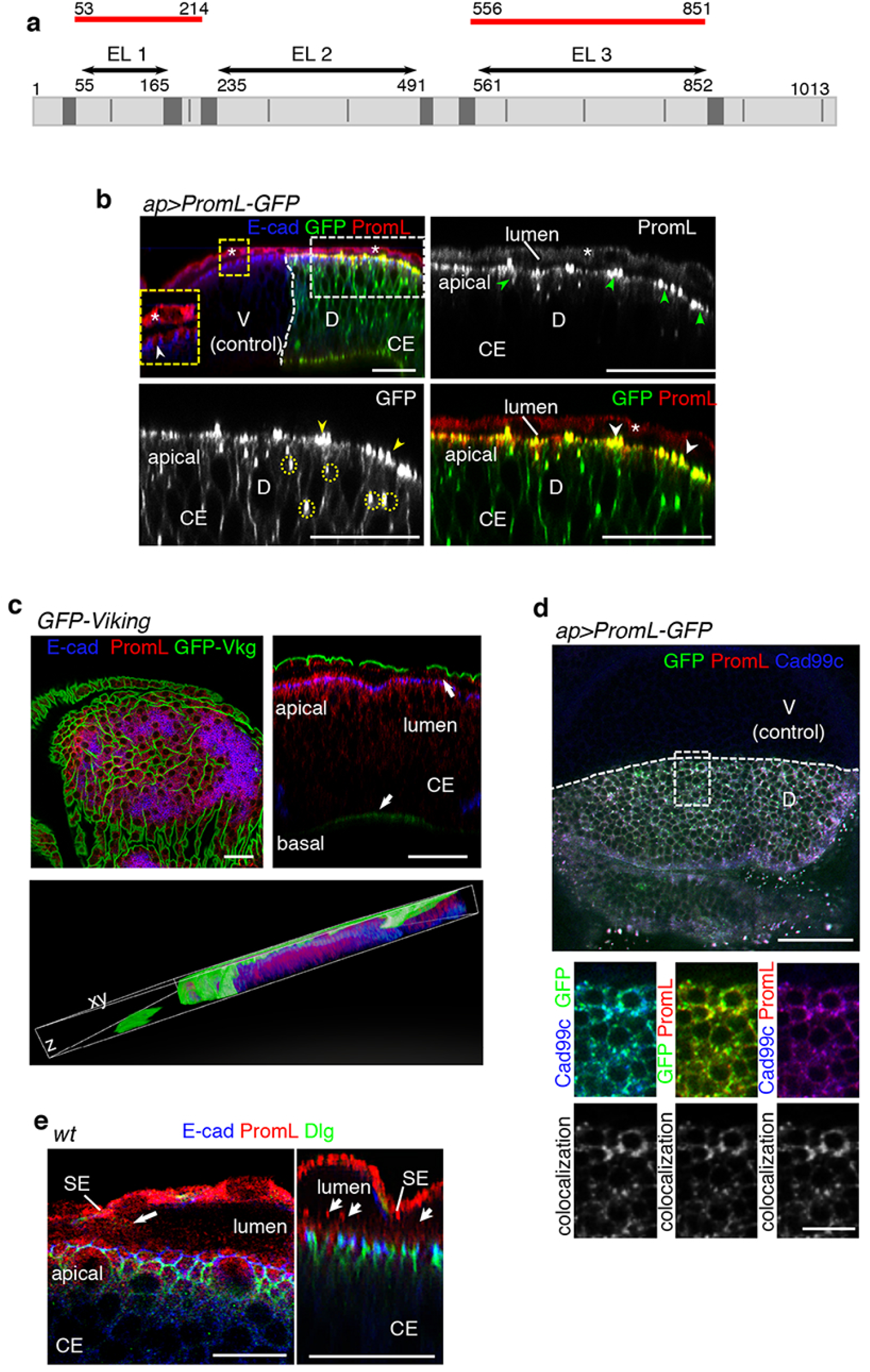

**Supplementary Figure S2.**
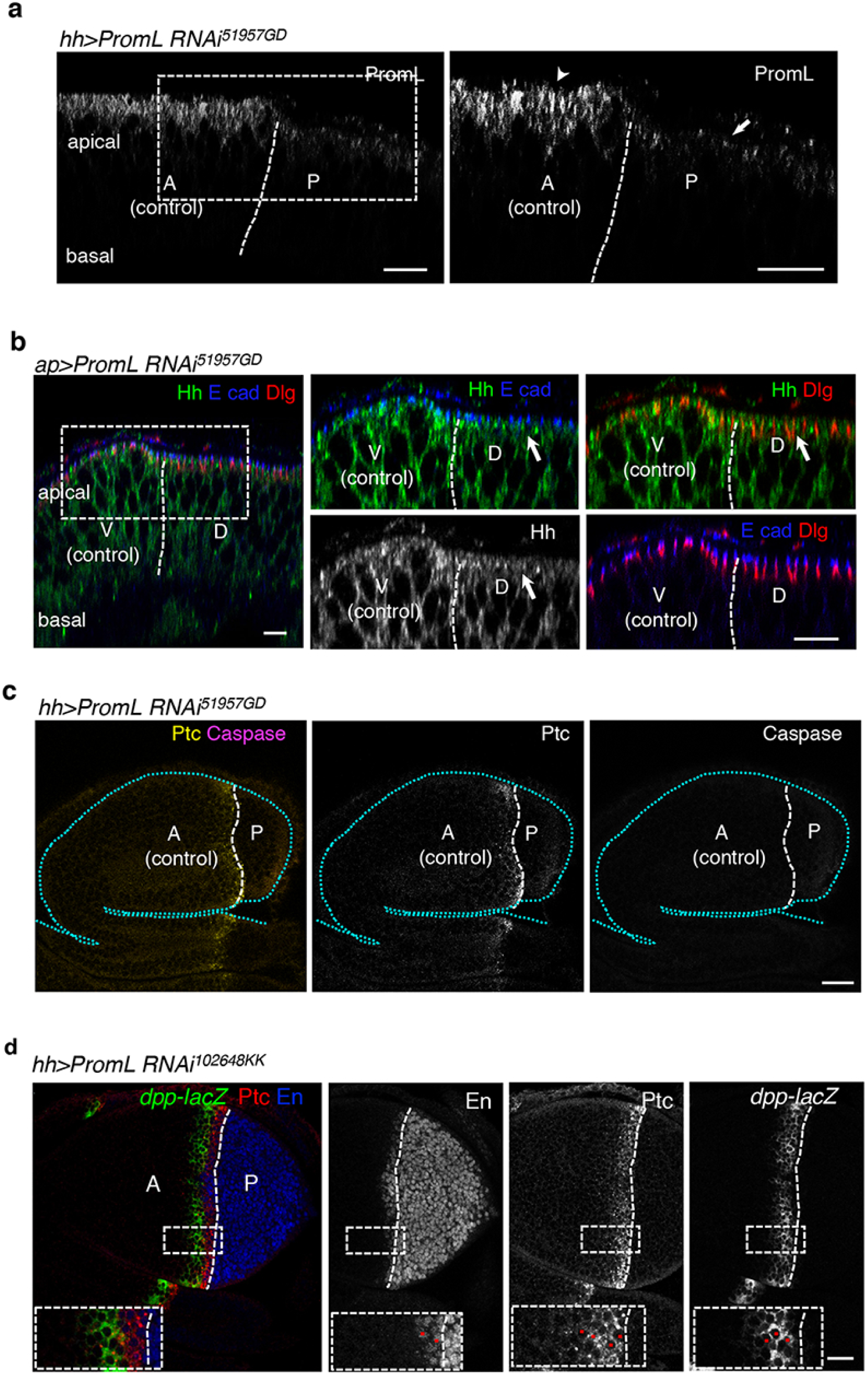

**Supplementary Figure S3.**
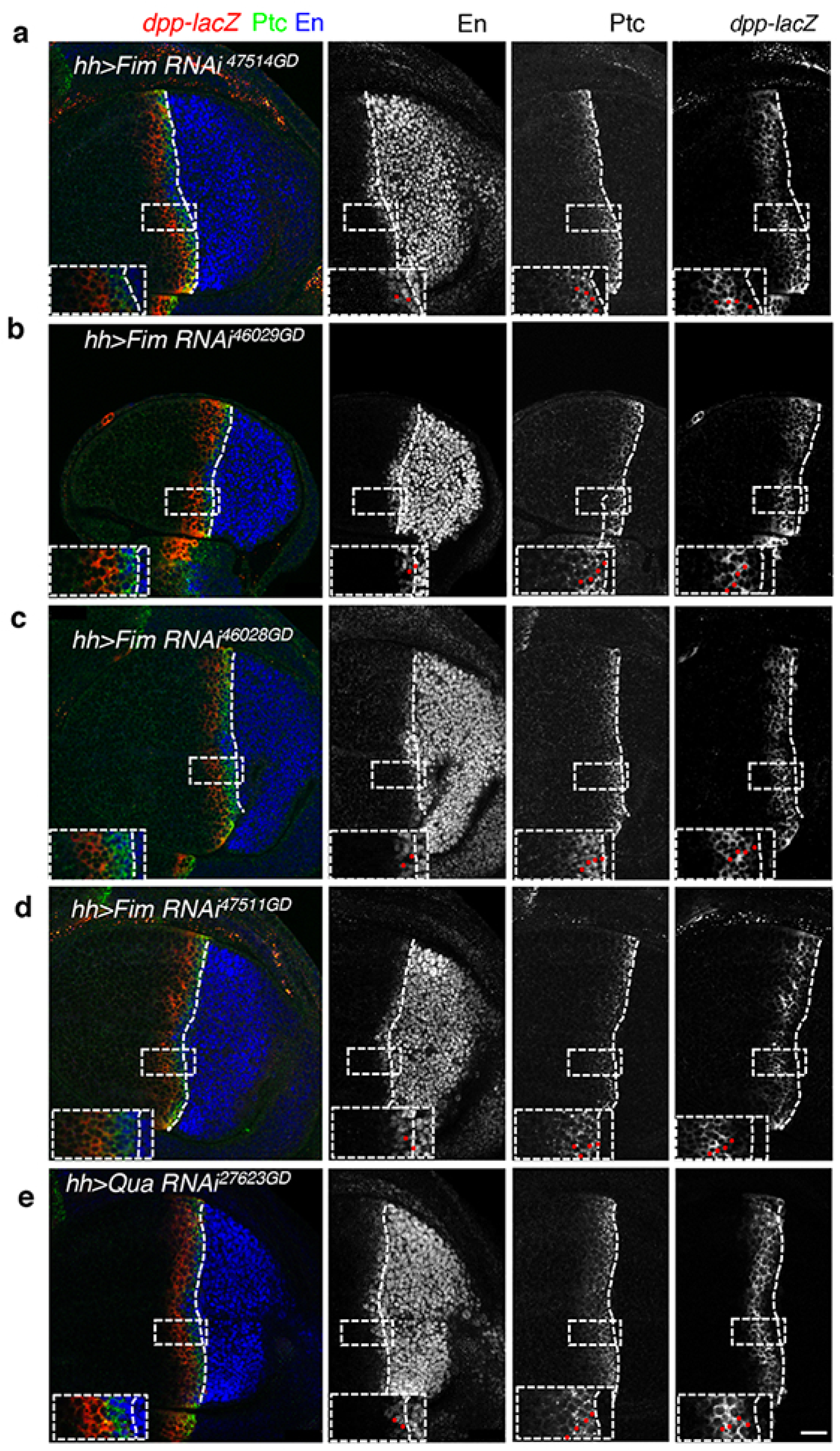

**Supplementary Figure S4.**
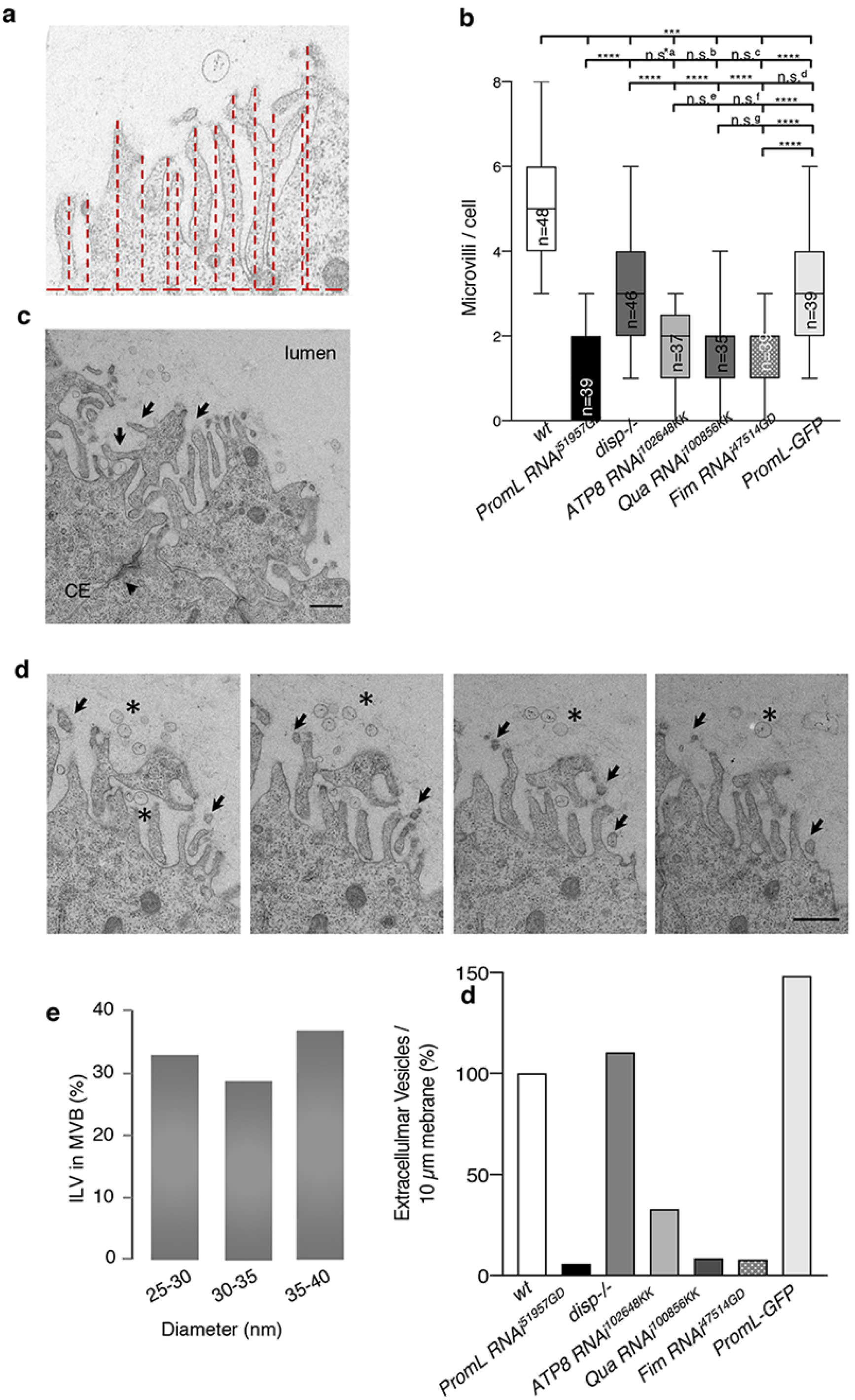

**Supplementary Figure S5.**
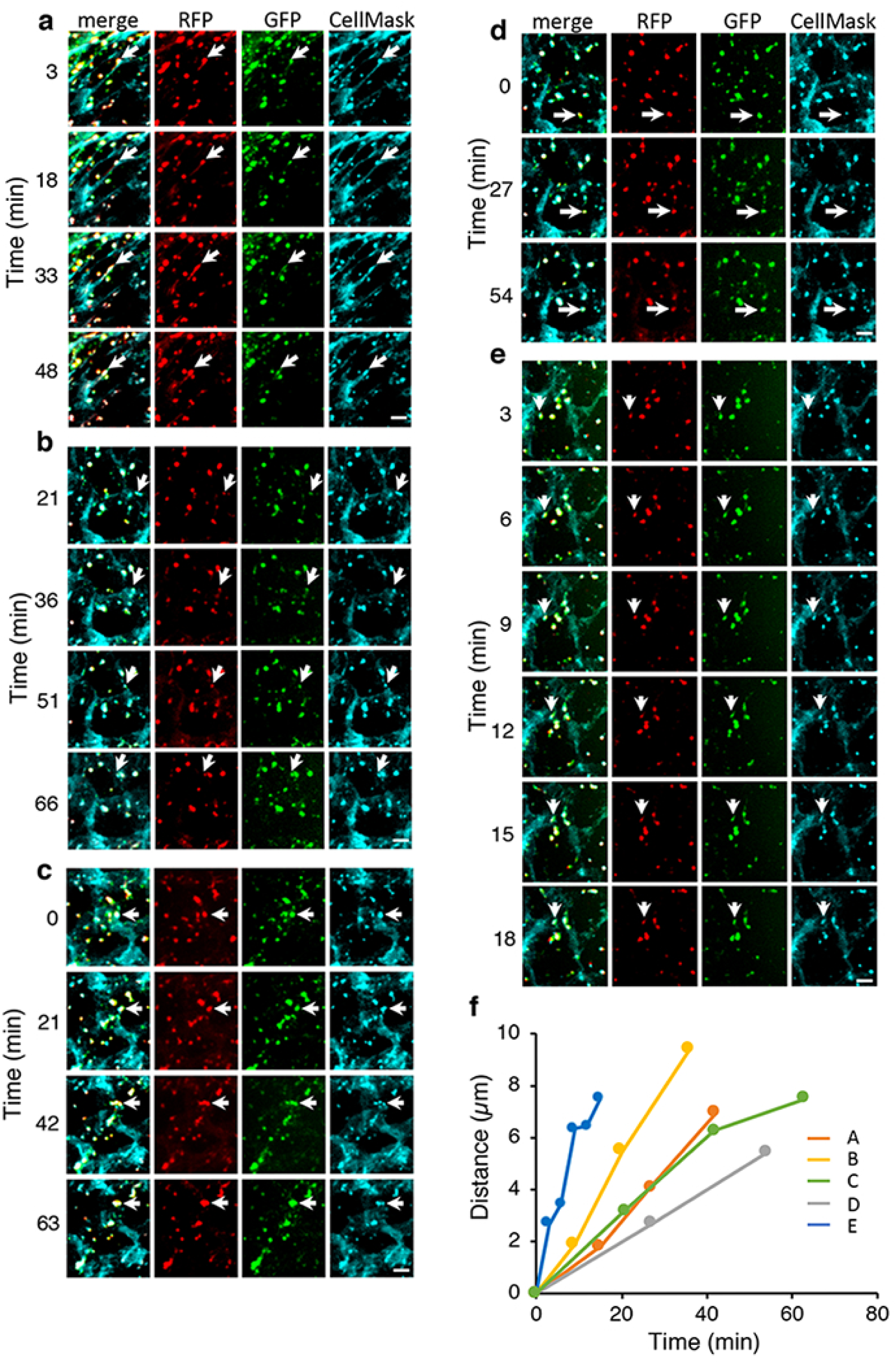

