## Supplementary info and Supplementary Figure Legends for "Microvilli-derived Extracellular Vesicles Govern Morphogenesis in *Drosophila* wing epithelium"

**This PDF file includes:**

- Methods
- Supplementary Figure Legends
- Captions for Movies 1-3
- References 29-32

**Other Supplementary Materials for this manuscript include the following:**

- Movies 1, 2, 3

### Methods

**Fly strains.** The following stocks were used: *$w^{1118}$*, *apGal4 (II), hhGal4 (III), UAS-Dicer2 (III)* from the Bloomington Stock Center and are listed in Flybase (www.flybase.org) and, protein-trap *$vkg^{G454}$* (*GFP-Vkg*)[16], *UAS-Hh-RFP* (II) from X. Lin (Cincinnati Children's Hospital Medical Center), *UAS-PromL-GFP (III)* from C. Dahmann (Dresden University of Technology. *$disp^{SO37707}$* is a null allele[20]. The UAS-RNAi lines against ATP8B ($ATP8B^{102648KK}$), PromL ($PromL^{102612KK}$, $PromL^{51957GD}$), Fimbrin ($Fim^{46028GD}$, $Fim^{46029GD}$, $Fim^{47511GD}$, $Fim^{47514GD}$ ), Quail ($Qua^{100856KK}$, $Qua^{27623GD}$) were from Vienna *Drosophila* Resource Center. Recombinant chromosomes were created for *a-Ga4l;UAS-Dicer2* and *UAS-Hh-RFP*; *UAS-PromL-GFP* by classical genetic techniques. All stocks were raised at 25 °C, unless otherwise mentioned.

**Antibody Generation.** cDNAs encoding two different regions (a.a. 53-214; a.a. 556-851) of the *D. melanogaster* PromL full length cDNA (UFO07496 clone; Drosophila Genomics Resource Center), were cloned into pStaby vector (Eurogentec) such as the peptides were tagged C-terminally with 6xHis. Both 6xHis peptides were expressed in E. coli using StabyExpress T7 kit, (Eurogentec) and purified on Ni-NTA resin (Invitrogen) by standard protocol. The peptides were soluble and injected individually into two guinea pigs (Eurogentec). To increase the antigenicity the obtained antisera were mixed, and the antisera were screened for antibodies against PromL using immunostaining of wing imaginal discs.

**Immunohistochemistry and fixed tissue imaging**. Third-instar wing imaginal discs were dissected and immunostained as previously described[5]. Briefly, the tissue was fixed in 4% paraformaldehyde for 20 minutes at room temperature, washed three times for 10 minutes with

PBS-T (PBS, 0.1% Triton X-100), and incubated with primary antibodies overnight at 4°C. After washing in PBS-T three times, discs were incubated with secondary antibody 1h at room temperature, and washed in PBS-T three times before mounting in Vectashield antifade mounting media (Vector Laboratories). Antibodies were used as follows:  mouse 4D9 monoclonal anti-En (Development Studies Hybridoma Bank University of Iowa – DSHB), 1:1,000; rabbit polyclonal anti-En (Santa Cruz), 1:1,000; mouse monoclonal anti-Ptc 1:400[29]; chicken polyclonal anti-βgal, 1:1,000 (Gen Tex); monoclonal affinity-purified rabbit “Calvados” polyclonal anti-Hh, 1:200[30]; mouse 4D4 monoclonal anti-Wg (DSHB); guinea pig anti-PromL, 1:150 (this study); rabbit polyclonal anti-aPKC (Santa Cruz), 1:500; rat DCAD2 polyclonal anti-DE-cadherin (DSHB), 1:50; mouse 4F3 monoclonal anti-Dlg (DSHB); mouse monoclonal anti-GFP (Roche), 1:1,000; rabbit polyclonal anti-Cad99c from D. Godt (University of Toronto), 1:1000; rabbit polyclonal anti-caspase 3, 1:500 (Cell signaling). Fluorescent secondary antibodies were used at 1:100 for Cy3-conjugated donkey anti-rat, Cy3- or Cy5-conjugated goat anti-mouse, Cy3- or Cy5-conjugated goat anti-rabbit, and Cy3-conjugated donkey anti-chicken (Jackson Laboratory), and 1:500 for 488 goat anti-mouse, anti-rat, and anti-rabbit (Life Technologies). Images were obtained with a Leica TCS SP5 or SP8 confocal microscope and processed with Image J. *XY* and *Z* sections are single sections captured at the appropriate level (apical, subapical, or lateral). Data analysis, curve fitting, and presentation were done with ImageJ, Excel and GraphPad Prim software.

**Transmission Electron Microscopy (TEM)**. Third-instar wing imaginal discs were dissected, fixed in 1.5% glutaraldehyde in 0.075 M cacodylate buffer (pH 7,2) overnight at 4°C, and post fixed in 1% osmium – 1.5% potassium ferrocyanide. The samples were then dehydrated in increasing concentrations of ethanol solutions, positioned perpendicularly (with the apical surface facing the top) and embedded in EPON. Ultrathin EPON sections (40-60 µm-thick sections depending on the genotype and the size of the wing discs), along the anterior/posterior axis were collected on formvar-coated slot grids and contrasted with uranyl acetate and lead citrate. Electron micrographs were acquired on a Tecnai Spirit transmission electron microscope (Tecnai Spirit, Thermo Fischer Scientific) equipped with 4k CCD camera (Quemesa, Soft Imaging System). Images were taken along the anterior/posterior axis enabling the visualization of the entire discs.

For Electron Tomography 300 nm thick sections were randomly labeled on the 2 sides with protein A-gold 15-nm and poststained with 2% uranylacetate. Tilt series (angular range from −60° to + 60° with 1° increments) were recorded by using EMtools (TVIPS) on a 200-kV transmission electron microscope (Tecnai 20 $LaB_6$, Thermo Fischer Scientific) and used for reconstructing tomograms. Projection images (2024 x 2024 pixels) were recorded with a CCD camera (Temcam F416; TVIPS). Alignment of the tilt series and tomogram computing (resolution-weighted back-projection) were carried out by using the eTomo (IMOD). The 15 nm gold particles at the surface of the sections were used as fiducial markers. Manual contouring of the tomograms was done by using the IMOD program[31] .

For Immunoelectron Microscopy (IEM), third-instar wing imaginal discs were fixed in a mixture of 2% PFA and 0,2 % glutaraldehyde in a 0,1M phosphate buffer pH 7,4 for 2h and processed for ultracryomicrotomy as described[32]. Ultrathin sections were prepared with an ultracryomicrotome UC7 (Leica), incubated with a rabbit anti-Hh antibody a gift from T. Kornberg (University of California, San Franscisco), with a rabbit anti-GFP, 1:120 (Molecular Probes),

**Live imaging.** Third-instar wing imaginal discs were dissected in Schneider's *Drosophila* Medium (21720-024, Gibco) supplemented with 10% heat-inactivated fetal bovine serum (10500, Gibco),

Penicillin (100 units ml$^{-1}$) and Streptomycin (100 µg ml$^{-1}$) (Penicillin-Streptomycin 15140, Gibco). Two to four wing discs were placed on a glass bottom 35 mm dish (P35G-1.5-14-C, MatTek Corporation) with 10 µl of medium containing CellMask (deep red plasma membrane stain; Molecular Probes) (1:1000), covered with a permeable membrane (Standard membrane kit, YSI), and sealed around the membrane borders with oil 10S Voltalef (VWR BDH Prolabo). Images were recorded using a Yokagawa CSU-W1 spinning head mounted on a Nikon TiE inverted microscope. The microscope was equipped with an EMCCD 1200 x1200 Prime95B (Photometrics) and controlled by the Metamorph software 7.8 (Molecular Devices). In most experiments, $z$-stacks at 1 µm intervals were acquired every 3 min using a 40 × NA 1.15 water immersion objective, but only the most in-focus planes were used for image analysis. Images were processed with ImageJ. All movies were analysed with Fiji and displayed at a rate of 7 frames per second.

**Manual tracking of extracellular vesicles.** EVs were tracked using the MTrackJ plugin of ImageJ. The tracking was performed in a summed $z$-projections taking into account 167 plans at 1 µm intervals and recorded every 3 min within a 25 µm x 30 µm delimited regions. Travel distances were analysed within the indicated time-window, measured using the line tool in ImageJ and then calculated using the following formula:

$$\text{distance } (ab) = \sqrt{(xb - xa)2 + (yb - ya)2}$$

where $(xa, ya)$ and $(xb, yb)$ are the coordinates of the initial and end points respectively.

**Quantification and Statistic analysis.** To quantify the colocalisation of PromL and Hh signals, the uppermost apical region of wing discs was delimited on single $z$-sections. For better accuracy, the intensity threshold for each channel has been applied manually. The colocalisation was computed as the mean of Hh-PromL intensity divided by the mean of Hh intensity alone. Data represent mean ± SEM.

EM micrographs of the different genotypes were used to manually quantify the number of microvilli and EVs observed. Since the microvilli were often tilted, we measured their length from their tip to the planar cell surface with an angle of 90°C (Supplementary Fig. 4a).

The diameter of EVs and ILVs was evaluated using iTEM software (Soft Imaging System, EMSIS GmbH, Germany). Data represent generally mean ± SEM.

The presence of Hh, PromL or both on EVs in A compartment was estimated using the algorithm described below. Given the positions of the $x_i$ particles clicked in the red channel, the algorithm defines the particle as a vesicle if the maximum intensity in the blue channel $I_b$, in the vicinity of the radius ($r$), is greater than a fixed threshold, such that:

$$\max_{B(x_i,R)} I_b > T_b$$ , where $B(x_i, R)$ is the ball of radius $r$ and center $x_i$.

The algorithm defines that the vesicle is also green on the same principle, based on the intensity on the green channel $I_g$, in the same vicinity:

$$\max_{B(x_i,R)} I\_g > T_g$$

We defined $r = 5$ pixels, $T_b = 800, T_g = 170$, and manual validations were performed to obtain more accurate results.

Data analysis, curve fitting, and presentation were done with ImageJ, Excel and GraphPad Prism software.

**Supplementary Figure 1**. **a,** Topology of the PromL protein. Dark grey boxes denote the predicted transmembrane domains. EL, extracellular loops. Red lines indicate the chosen sequences for the antibody generation. **b-e,** Confocal single sections of wing imaginal disc of the indicated genotypes labeled with anti-GFP (green; white), PromL (red; white), E-cad (blue), Cad99c (blue), and Dlg (green). **b**,**d**, The wild-type ventral (V) compartment, allows the comparison between both compartments within the same disc. **b**, PromL distributed to the squamous epithelium (asterisk), and to the apical surface of columnar cells (CE) (green arrowheads) above the subapical marker E-cad (white arrowhead). Bottom left: yellow arrows and dotted circles mark the distribution of PromL-GFP to the apical and basolateral domains respectively. Bottom right shows examples of colocalisation between PromL-GFP and endogenous PromL (white arrowheads). **c**, Right: PromL distributes above E-cad and below endogenous Viking protein which labels the basal membrane (arrows). Bottom: 3D volumetric reconstruction (19 plans, spaced 1 µm apart). **d**, PromL, Cad99c and PromL-GFP colocalise at the apical (high magnification of boxed region). Brightness and contrast were increased for a better visualization. **e**, PromL distributed above E-cad and Dlg. The dot-like pattern of PromL (white arrow) at the interface between SE and CE reflects the pseudostratified array of columnar cells, and is compatible with the organization of microvilli at the apical plasma membrane. Dotted lines in **b**,**d** delimite V and D compartments. Scale bars: 20 µm; 5 µm in magnified images in **d**. Images shown in **b** and **e** are representative of n=5 independent experiments. Images shown in **c** and **d** are representative of n=3 independent experiments.

**Suplementary Figure 2**. **a-d**, Confocal images of discs of the indicated genotypes stained for PromL (white), Hh (green; white), E-cad (blue), Dlg (red), Caspase III (magenta; white), Ptc (yellow; red; white), En (blue; white) and for β-gal (reflecting the expression of the reporter gene *dpp-lacZ*; red; white). **a**, Righ: shows the specific dot-like pattern of PromL in A control cells (arrowhead), and the absence of PromL staining in posterior (P) depleted cells (arrow). **b**, In D cells, Hh distributed at the basolateral, below E-cad, at the level of Dlg (white arrows). The distribution of E-cad and Dlg is not affected upon PromL depletion (compare V/D compartment). **c**, Note that apoptosis levels were not increased upon PromL depletion, indicating and absence of tissue stress. **d**, Depletion of PromL triggers a decrease in the range of cells expressing *dpp*, reducing it to 3–4 cells. The short-range targets En and Ptc are not changed. Red dots in the inserts depict the number of cells (see Table 1). Broken lines delimit the A/P compartments. Scale bars: 20 μm.

**Supplementary Figure 3**. **a-e**, XY single confocal sections of discs of the indicated genotypes stained for En (blue; white), Ptc (green; white), and β-gal (red; white). Insets depict the number of cells expressing En, Ptc and *dpp-lacZ.* Note the decrease in the range of cells expressing *dpp* in all genotypes as compared to *wt* (Fig. 1j and Table 1). No changes are observed for En and Ptc. Red dots in the insects indicate the number of cells. Broken lines delimit the A/P compartments. Scale bars: 20 μm. Scale bars: 20 μm.

**Supplementary Figure 4. a,** The length of microvilli was measured by drawing a perpendicular line from their tip to the planar cell surface with an angle of 90° as illustrated in the cartoon. **b**, Graph showing the number of microvilli / cell for the indicated genotypes. Mean ± SEM analyzed by unpaired *t* test **** $p < 0.0001$, n.s., not significant; from n.s[a] -n.s.[g]. : 0.0269; 0.0812; 0.1050; 0.2724; 0.5647; 0.4023; 0.8096. The number of cells counted for each genotype is indicated in each column bars. **c**, **d**, TEM micrograph of a *wt* wing disc showing (**c**) a transverse or (**d**) serial sections of 65 nm thickness (260 nm total). **c**, Arrows point to microvilli, arrowheads to basolateral junctions. **d**, Asterisk indicate free EVs within the lumen. Note that EVs are less electron dense than fragmented microvilli (arrows). Scale bar: 200 nm. **e**, Distribution of ILVs based on their diameter (n=38). Note that the diameter of ILV is smaller than the one of EVs (Fig. 4b). **f**, Histogram of the percentage of EVs detected for 10 µm of the apical membrane for the indicated genotypes.

**Supplementary Figure 5. a-e,** Frames from of *hh>Hh-RFP; PromL-GFP* discs imaged at 1 µm intervals every 3 min, showing additional examples of Hh/PromL-EV movement in the anterior compartment. PromL-GFP (green), Hh-RFP (red) and plasma membrane (Cell Mask; Cyan). Arrows depict the EVs. Scale bar: 5 µm. **f**, The distances traveled by Hh/PromL-EV from panels **a-e** were linearly fitted. Note that the distance gradually increased with time.

**Movie 1**.**Tomographic reconstruction and the 3D model of the apical plasma membrane of wing imaginal disc epithelium**. Buds (red) emanate from the microvillar membrane (yellow) and remain connected or are released into the lumen, confirming that microvilli are the sites of EV formation.

**Movie 2. Virtual sections though the 3D reconstruction of a *wt* disc.** EVs circled in blue show no contact with microvilli. EVs circled in red are attached to microvilli membrane. The microvilli membranes are outlined in orange

**Movie 3**. **Hh/PromL-rich EVs emanating from posterior cells travel to and in the anterior compartment**. *hh>Hh-RFP; PromL-GFP* live discs were imaged at 1 µm intervals every 3 min for 90 min. PromL-GFP (green), Hh-RFP (red), epithelial cell membrane (Cyan). Scale bar : 5 µm.
